# Protective Transfer: Maternal passive immunization with a rotavirus-neutralizing dimeric IgA protects against rotavirus disease in suckling neonates

**DOI:** 10.1101/2021.09.21.461116

**Authors:** SN Langel, JT Steppe, J Chang, T Travieso, H Webster, CE Otero, LE Williamson, JE Crowe, HB Greenberg, H Wu, C Hornik, K Mansouri, RJ Edwards, V Stalls, P Acharya, M Blasi, SR Permar

## Abstract

Breast milk secretory IgA antibodies provide a first line of defense against enteric infections. Despite this and an effective vaccine, human rotaviruses (RVs) remain the leading cause of severe infectious diarrhea in children in low- and middle-income countries (LMIC) where vaccine efficacy is lower than that of developed nations. Therapeutic strategies that deliver potently neutralizing antibodies into milk could provide protection against enteric pathogens such as RVs. We developed a murine model of maternal protective-transfer using systemic administration of a dimeric IgA (dIgA) monoclonal antibody. We confirmed that systemically-administered dIgA passively transferred into milk and stomach of suckling pups in a dose-dependent manner. We then demonstrated that systemic administration of an engineered potent RV-neutralizing dIgA (mAb41) in lactating dams protected suckling pups from RV-induced diarrhea. This maternal protective-transfer immunization platform could be an effective strategy to improve infant mortality against enteric infections, particularly in LMIC with high rates of breastfeeding.

**GRAPHICAL ABSTRACT:** 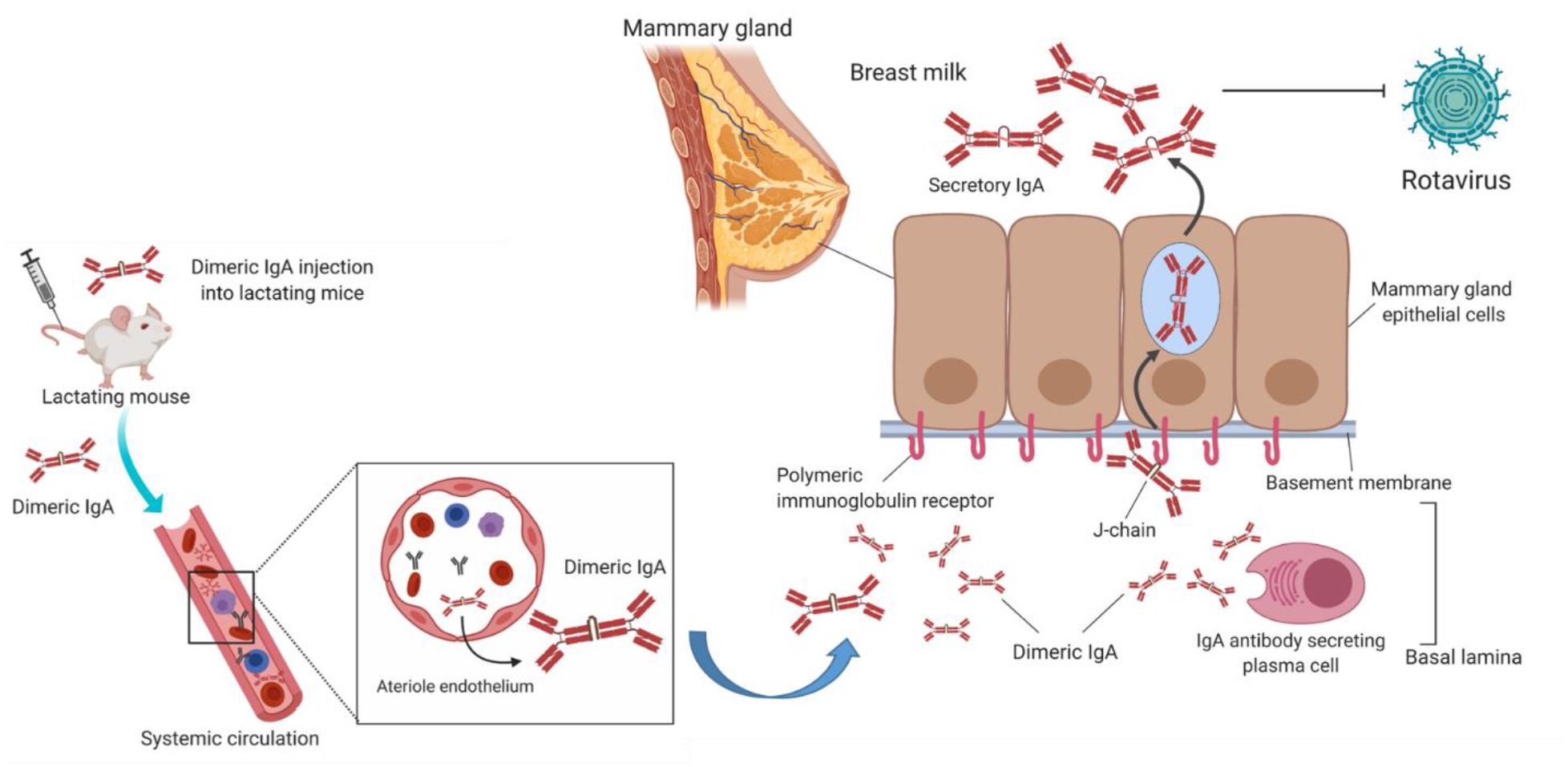

## INTRODUCTION

Rotavirus (RV), a common enteric pathogen, is responsible for ∼125,000-215,000 deaths in children <5 years and is the leading cause of gastroenteris-related hospitalizations worldwide, particularly in low and middle-income countries (LMIC) (Burke et al., 2019). While oral live, attenuated RV vaccines have significantly reduced RV-associated disease and death worldwide (Burnett et al., 2017), RV vaccines demonstrate lower efficacy in children in LMIC (40-60%) compared to those in high-income countries (80-90%) (Jonesteller et al., 2017; Mwila et al., 2017). The combination of decreased RV vaccine efficacy, high rates of RV exposure and an immature immune system creates a ‘window of susceptibility’ when infants and toddlers are particularly vulnerable. Strategies to provide additional antibody-mediated protection are needed to narrow this critical window of vulnerability and further decrease RV-associated disease and death.

While maternal immunization could be employed as a strategy to boost protective antibodies in breast milk, pregnant women in areas with high RV-associated morbidity and mortality experience undernutrition (Desyibelew and Dadi, 2019), micronutrient deficiencies (Harika et al., 2017) and chronic enteropathies that may impact generation of potently neutralizing anti-RV responses. Indeed, previous studies have demonstrated that socioeconomic status plays a role in generation of anti-RV antibodies (Ray et al., 2007; Trang et al., 2014). Additionally, maternal vaccination may lead to maternal antibody interference and decreased immunogenicity in vaccinated infants (Appaiahgari et al., 2014; Otero et al., 2020). Based on this, other therapeutic strategies for delivery of potently RV-neutralizing antibodies are needed.

In humans and animal models, a high titer of intestinal RV-specific IgA is a correlate of protection against RV infection and illness (Blutt et al., 2012; Matson et al., 1993; Tô et al., 1998). Therefore, an ideal way to increase protection against RV infection is to provide RV-neutralizing IgA to the infant gut via breast milk. Breast milk contains mostly secretory IgA (sIgA) antibodies; proteolytically stable dimers connected by a joining (J) chain (Corthesy, 2013; Hurley and Theil, 2011). Passive transfer of sIgA into breast milk depends on locally-produced dimeric IgA (dIgA) binding to polymeric immunoglobulin receptor (pIgR) on the basal side of mammary gland epithelial cells via the J-chain (Goldblum et al., 1975; Johansen et al., 1999; Tuaillon et al., 2009). The dIgA-pIgR complex is then transferred across mammary gland epithelium and secreted into breast milk as sIgA (De Groot et al., 2000). Once in milk, sIgA provides immune protection by neutralizing enteric toxins and pathogenic microorganisms and mediating microbiota colonization through exclusion of exogenous competitors (Mantis et al., 2011; Pabst and Slack, 2020). In LMIC, high RV-specific antibody titers in breast milk were associated with decreased incidence of infant RV diarrhea, and partial breastfeeding significantly increased the risk of infant mortality due to diarrheal disease compared to exclusive breastfeeding (Arifeen et al., 2001; Jayashree et al., 1988). However, milk RV-neutralizing antibody titers vary greatly in women from LMIC and may not provide adequate levels of protection (Trang et al., 2014). Therefore, developing dIgA therapeutics that are designed for passive transfer into mucosal compartments, including breast milk, is a novel strategy for enhancing protection of the mother-infant dyad against infectious enteric pathogens.

Here, we developed a murine model of maternal systemic passive antibody immunization for transfer into breast milk using a murine dIgA monoclonal antibody (mAb). We demonstrate that following systemic administration of a dIgA mAb in lactating mice, the mAb is rapidly transported into milk and quickly detected in pup stomach content in a dose-dependent manner. We then engineered a murine dIgA version of a human RV-neutralizing IgG (mAb41) (Nair et al., 2017) and optimized it for enhanced production of dIgA antibodies *in vitro*. We show that dams systemically injected with the RV-neutralizing dIgA mAb provided protection to their pups against RV-associated diarrhea. Our results demonstrate that dIgA can passively transfer out of circulation and into milk to provide a novel strategy for protection against RV disease in suckling neonates. These studies support the need for clinical assessment of maternal protective transfer via passive immunization with dIgA, particularly in LMIC where infant enteric disease burden is high.

## RESULTS

### Maternal systemic administration of dIgA results in antibody transfer into milk of lactating mice and the gastrointestinal tract of their pups

To determine whether systemically-administered dIgA passively transfers into milk, we first used a dIgA mAb generated from a previously-described non-neutralizing RV-specific murine dIgA 7D9 hybridoma cell line (Burns et al., 1996). We confirmed 7D9 dIgA binding to cognate antigen, the RV capsid protein VP6, and the J-chain receptor pIgR (Figure 1A) demonstrating that 7D9 contains the J-chain needed for pIgR-mediated passive transfer into milk (Figure 1A). The purified 7D9 hybridoma supernatant was also confirmed to contain dimeric antibodies by negative stain electron microscopy (NSEM) (Figure 1B) and size-exclusion chromatography (SEC) (Figure S1A). While dimers were the most prevalent antibody species present, we identified small contributions from higher order IgA species, like tetrameric IgA (Figure S1B).

**Figure 1.**
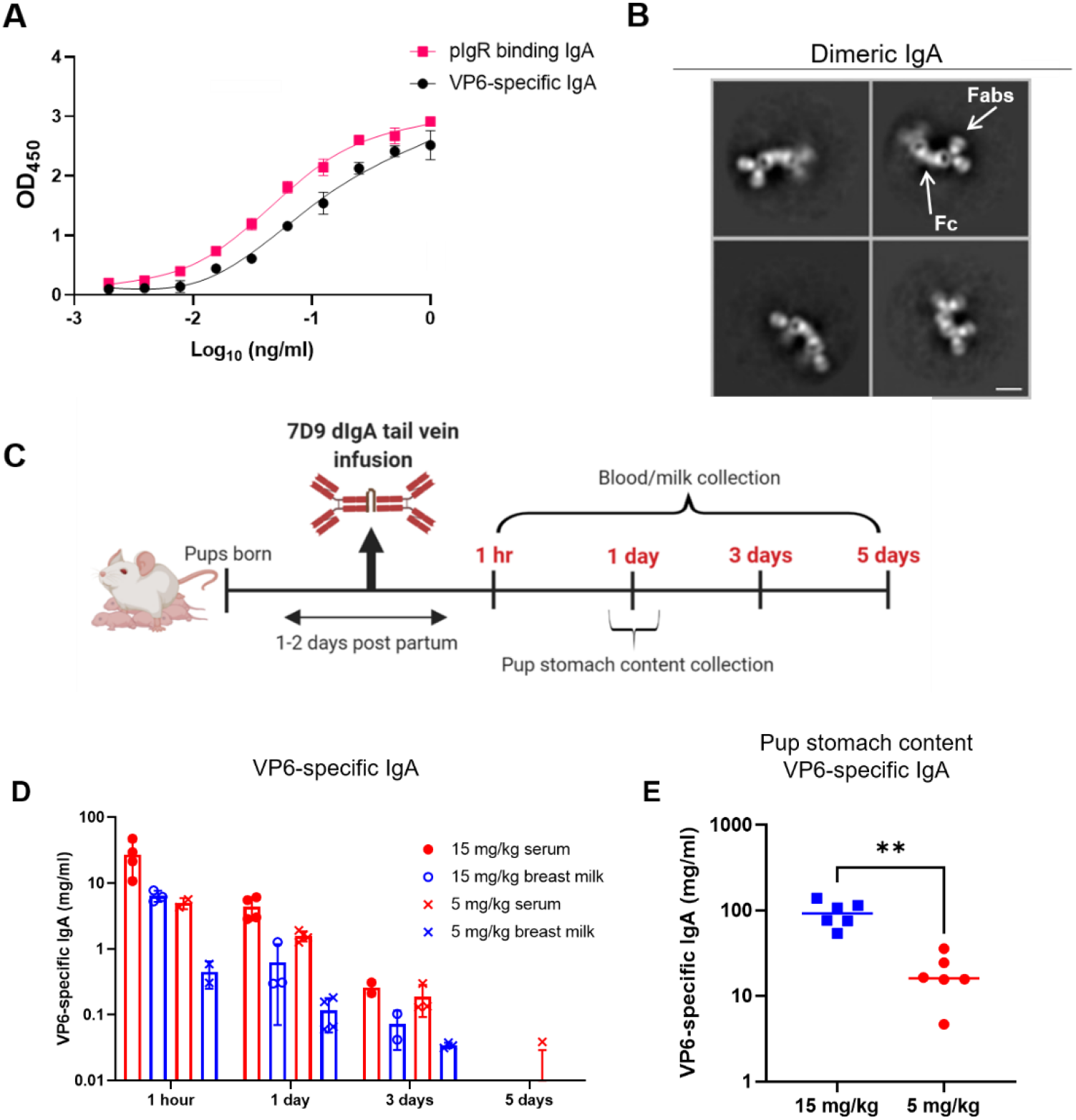
Dimeric IgA antibodies administered systemically to lactating mice are transferred to milk and the stomach contents of suckling pups. (A) The rotavirus (RV) VP6-specific, non-neutralizing 7D9 hybridoma derived antibodies bound to RV VP6 (black circles) and polymeric immunoglobulin receptor (pIgR; magenta squares) via ELISA. Data are plotted as mean ± SD of replicates. (B) Negative stain electron microscopy representative images of the hybridoma-purified 7D9 dimeric IgA (dIgA) antibodies. The Fab and Fc regions are indicated respectively. Scale bar represents 10 nm. (C) Schematic of tail vein injections of BALB/c lactating dams given 5 mg/kg or 15 mg/kg 7D9 dIgA at 1 to 2 days postpartum. Blood and milk were collected from dams at 1 hr and 1, 3 and 5 days post injection. A subset of pups (n=6) per treatment group were sacrificed at 1-day post injection to collect their stomach content. The schematic was created with Biorender. (D) 7D9 antibodies were detected in blood and milk of injected dams at 1 hr and 1, 3 and 5 days post injection via a RV VP6-specific IgA antibody ELISA. The 5 mg/kg (x) and 15 mg/kg (triangle) treatment groups are indicated for serum (red) and milk (blue). Data are plotted as mean ± SD and represent individual mice. (E) 7D9 antibodies were detected in the stomach content of suckling pups via a RV VP6-specific IgA antibody ELISA in a dose-dependent manner (5 mg/kg = red circles; 15 mg/kg = blue squares). Data are plotted as mean ± SD and represent individual pups. A significant difference between the compared groups (**p < 0.01) was determined using a Mann-Whitney U test.

To confirm passive transfer of 7D9 dIgA into milk, lactating BALB/c dams were infused with 5 mg/kg or 15 mg/kg of 7D9 at 1 to 2 days postpartum and plasma and milk were collected at 1 hr and 1, 3 and 5 days after infusion to assess antibody concentrations (Figure 1C). Peak plasma and milk concentrations of 7D9 antibodies, as measured by a VP6-specific IgA ELISA, were observed 1 hr after infusion in both the 5 mg/kg and 15 mg/kg groups (Figure 1D). To confirm transfer of 7D9 dIgA from milk to suckling pups, a subset of pups were sacrificed 1-day post infusion from both the 5 mg/kg (n=6) and 15 mg/kg (n=6) litters and their ntestinal contents were collected and analyzed for the presence of 7D9. 7D9 was detected in the stomach content of litters born to dams from both 5 mg/kg and 15 mg/kg groups in a dose-dependent manner (Figure 1E). 7D9 antibody levels precipitously dropped 1-day post infusion in both the serum and milk of lactating dams. Low or undetectable levels were observed by day 5 post infusion. This kinetics suggest rapid transfer of 7D9 dIgA to mucosal secretions, including milk. Indeed, 7D9 IgA was also detected in intestinal and rectal content, at low levels in saliva but not in vaginal washes (Figure S2A, B), which suggests pIgR expression differences at different mucosal sites. These data demonstrate systemically-administered dIgA is efficiently transferred from the systemic circulation to milk and other mucosal compartments of mouse dams and subsequently into pup stomach content.

### Engineering and recombinant production of a RV-neutralizing mouse-human chimeric dIgA

After demonstrating that systemically-administered 7D9 dIgA can passively transfer from the periphery to milk using our murine lactating mouse model, we aimed to engineer a RV-neutralizing dIgA that could provide protection against RV-induced diarrhea. We constructed a dIgA version of a previously-isolated (Nair et al., 2017) potently-neutralizing RV VP4-specific mAb (mAb#41 or mAb41) by replacing the human IgG1 with the murine IgA constant region and adding the BALB/c J-chain gene on the same open reading frame (Figure 2A). Characterization of the recombinantly produced mAb41 dIgA by ELISA, demonstrated that not all the produced antibody bound to pIgR (Figure 2B), suggesting that in addition to dIgA, non-dimeric or aggregated IgA species were also present. Indeed, NSEM (Figure 3A) and SEC (Figure 3B) revealed multiple IgA species including monomeric and dimeric (Figure 3A) and aggregated (Figure S3) antibodies. We next fractionated the different IgA species based on size and evaluated them for their ability to bind (Figure 3C) and neutralize (Figure 3D) RV. Interestingly, the fraction suspected to be enriched in dIgA (fraction 29-39) had the greatest RV-binding and neutralization capacity compared to the other isolated fractions.

**Figure 2.**
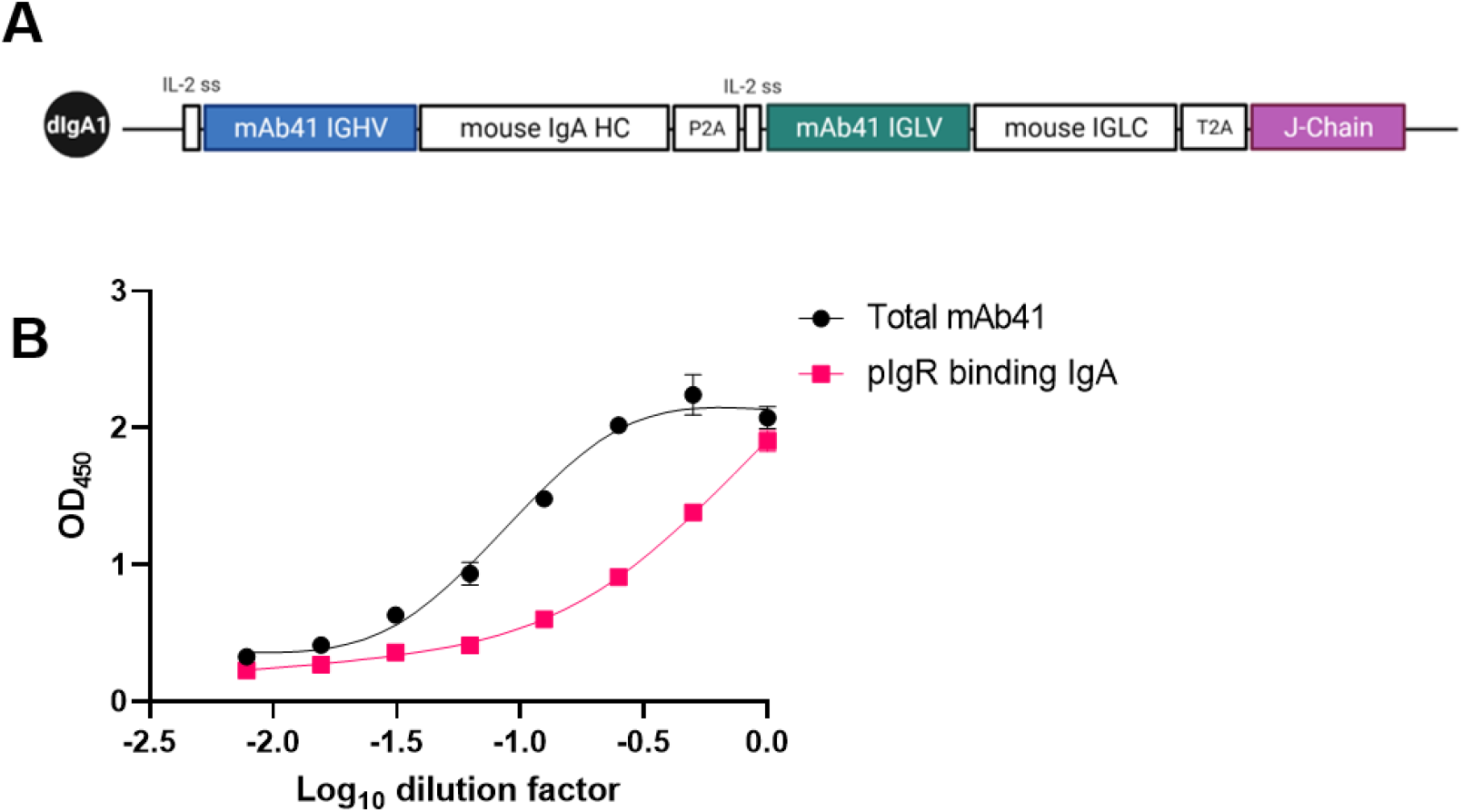
Recombinant production of an RV-neutralizing mouse-human chimeric dIgA antibody. (A) Schematic of the plasmid used to produce the mouse-human chimeric mAb41 dimeric IgA. The mAb41 immuoglobulin heavy chain variable (IGHV) gene (blue box), immunoglobulin light chain variable (IGLV) gene (green box), and the joining chain (J-chain) gene (purple box), are indicated in that order. Schematic created with Biorender. (B) J-chain containing IgA antibodies (mAb 41 = black circles) were detected via a polymeric immunoglobulin receptor (pIgR) binding ELISA. A positive control mAb (magenta squares) was included. Data are plotted as mean ± SD of replicates.

**Figure 3.**
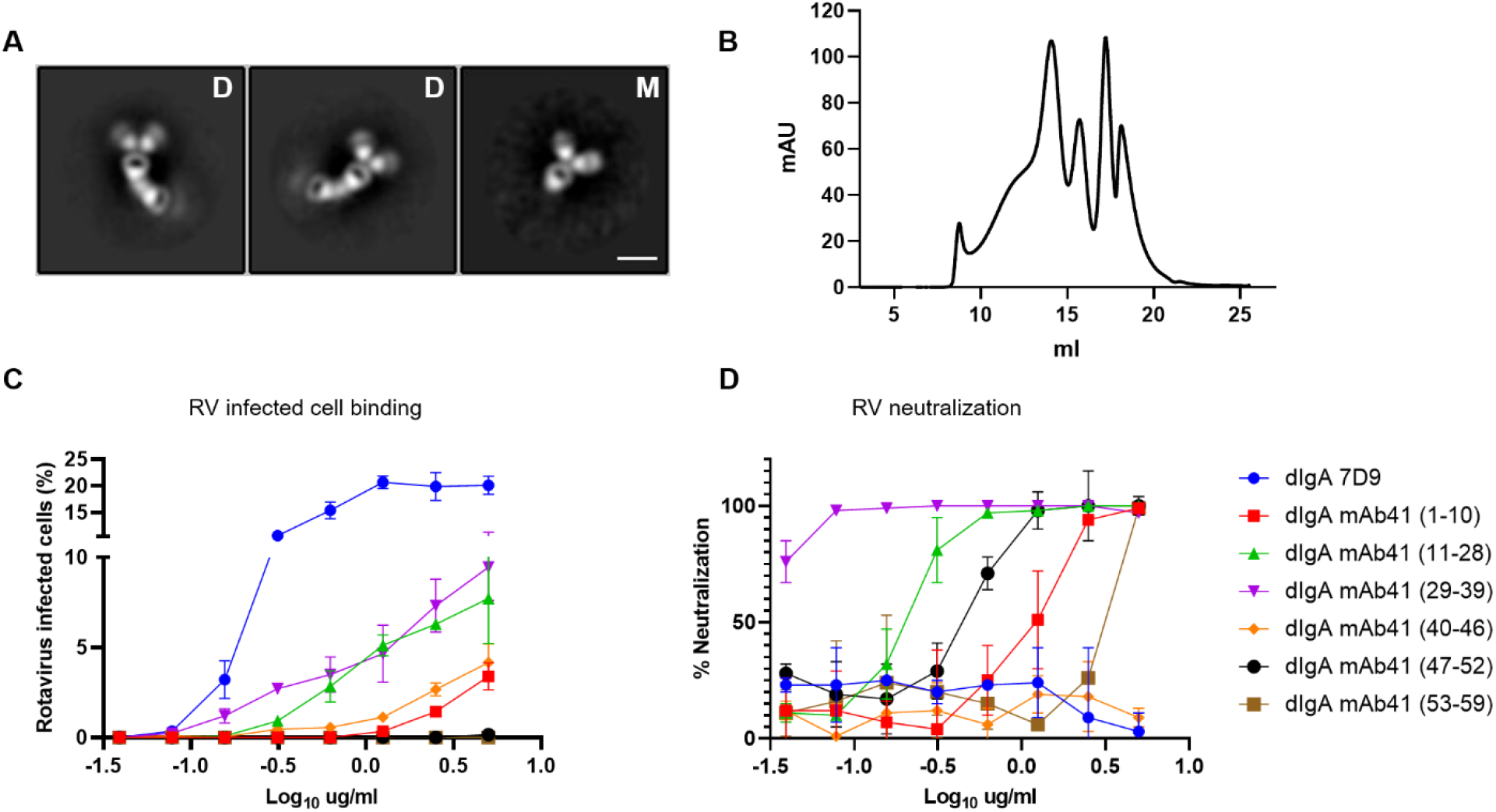
Characterization of recombinant mAb41 IgA antibodies. (A) Representative negative stain electron microscopy images of purified dimeric (D) and monomeric (M) mAb41 IgA antibodies. (B) Size exclusion chromatography using a Superose 6 10/300 GL revealed multiple different peaks. Each of the peaks were fractionated and the corresponding fractions (1-10, 11-28, 29-39, 40-46, 47-52 and 52-59) functionally characterized by rotavirus (RV)-infected cell binding (C) and neutralization assays (D). RV-infected MA104 cell binding assays revealed that fraction 29-39 bound the strongest to infected cells (C). Similarly, this fraction had the most potent neutralization of RV as determined via RV neutralization assays (D). Based on molecular mass this fraction contains dIgA. The non-neutralizing 7D9 dIgA (blue line) was used as a positive control for binding and a negative control for neutralization. Data are plotted as mean ± SD of replicates.

To generate a more homogenous product and skew antibody production towards increased dimer formation, we designed three additional dIgA constructs, where furin cleavage sites were added before the 2A self-cleaving peptides to enhance cleavage, and the J-chain was placed either in the middle (dIgA.2), at the end (dIgA.3) or at the beginning (dIgA.4) of the ORF (Figure 4A). All constructs produced large amounts of total mAb41 IgA antibodies (Figure 4B), however placement of the J-chain gene at the end of the construct (dIgA.1) resulted in the highest amount of pIgR binding IgA antibodies (Figure 4C).

**Figure 4.**
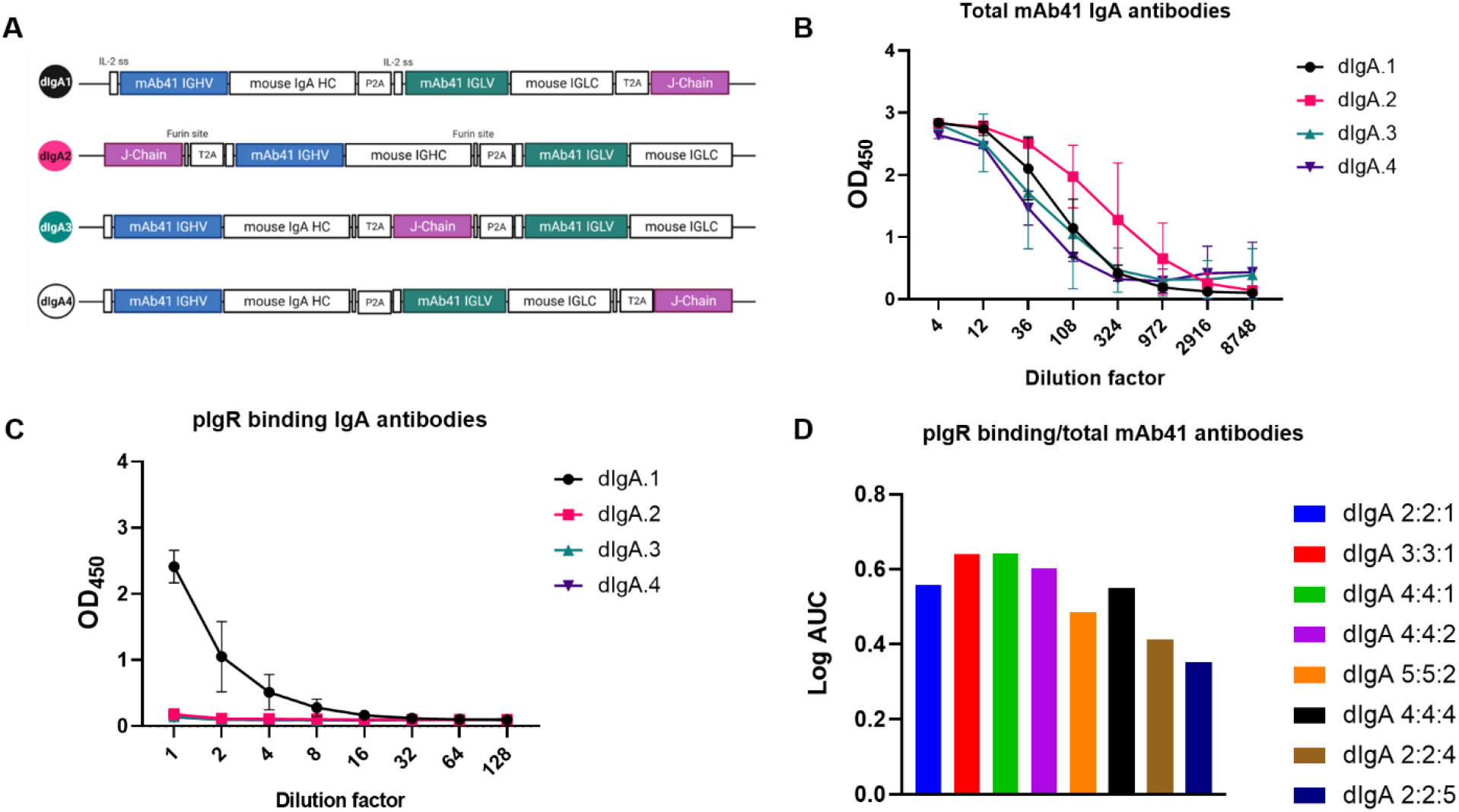
Antibody chains position and ratio impacts the recombinant production of RV-neutralizing mouse-human chimeric mAb41 dimeric IgA. (A) Schematic of the different constructs generated to determine if the position of the J-chain gene in the plasmid cassette impacts dimeric IgA (dIgA) production. Illustration created with Biorender. (B) All constructs produced high amounts of mAb41 IgA. However, placement of the J-chain gene at the end of the plasmid cassette (dIgA.1) resulted in the highest amount of pIgR binding IgA antibodies as determined by ELISA (C). Data are plotted as mean ± SD of experimental duplicates. (D) To determine the optimal ratio of heavy light and J-chain for dimeric IgA production, we co-transfected three separate plasmids each expressing either the heavy (H), the light (L) or the J-chain (J) in varying amounts as depicted on the graph. pIgR IgA antibodies and total mAb41 IgA antibodies were detected via ELISA. Log area under the curve (AUC) was calculated for each co-transfection and graphed as a ratio of pIgR binding IgA antibodies over total mAb41 IgA antibodies. Greater production of pIgR binding IgA antibodies was achieved when lower amounts of J-chain expressing plasmid were used compared to heavy and light chain plasmids. Data are plotted as mean ± SD of experimental duplicates.

To address whether modulating the ratio between the J-chain and the heavy and light chains resulted in increased dimer formation, we generated three additional plasmids each separately expressing the heavy [H], the light [L] or the J [J] chain genes and compared dimer production following cells transfection with increasing amounts of J-chain plasmid. Increasing the amount of J-chain DNA resulted in lower levels of pIgR-binding IgA (Figure 4 D), which is concordant with a previous report of recombinant IgA production (Lombana et al., 2019). The 4:4:1, 3:3:1 and 4:4:2 ratio of H:L:J chain plasmids produced the highest level of pIgR-binding dimers (Figure 4D).

### Recombinant mAb41 dIgA demonstrates greater RV-binding and neutralization potency than mAb41 monomeric IgA or IgG1 antibodies

To exclude functional contributions from aggregated dimeric/polymeric or monomeric IgA (mIgA) antibodies, recombinantly produced mAb41 dIgA was fractionated by SEC (Figure 5A) and the average molecular weight of 300 kDa was selected as fractionated mAb41 dIgA (f-dIgA mAb41), which is consistent with the expected mass of dIgA at approximately 335 kDa. NSEM demonstrated that the f-dIgA mAb41product contained only dimeric antibodies (Figure 5B). The f-mAb41 dIgA product was then tested for functional capacity in comparison with mIgA mAb41 and IgG mAb41. f-dIgA mAb41 demonstrated greater RV-binding capacity and neutralization activity (AUC=31.9; IC_90_ = 7.1 ng/ml) compared to mIgA (AUC=24.8; IC_90_ = 44.8 ng/ml) or IgG (AUC=19.7; IC_90_ = 33.01 ng/ml) (Figures 5C and 5D). This difference in neutralization activity was even greater when IC_90_ values were normalized for antibody molecular weight, resulting in a 42-fold reduction in IC_90_ value for f-dIgA mAb41 (21.2 pM) compared to mIgA (298.3 pM) and IgG (220.5 pM).

**Figure 5.**
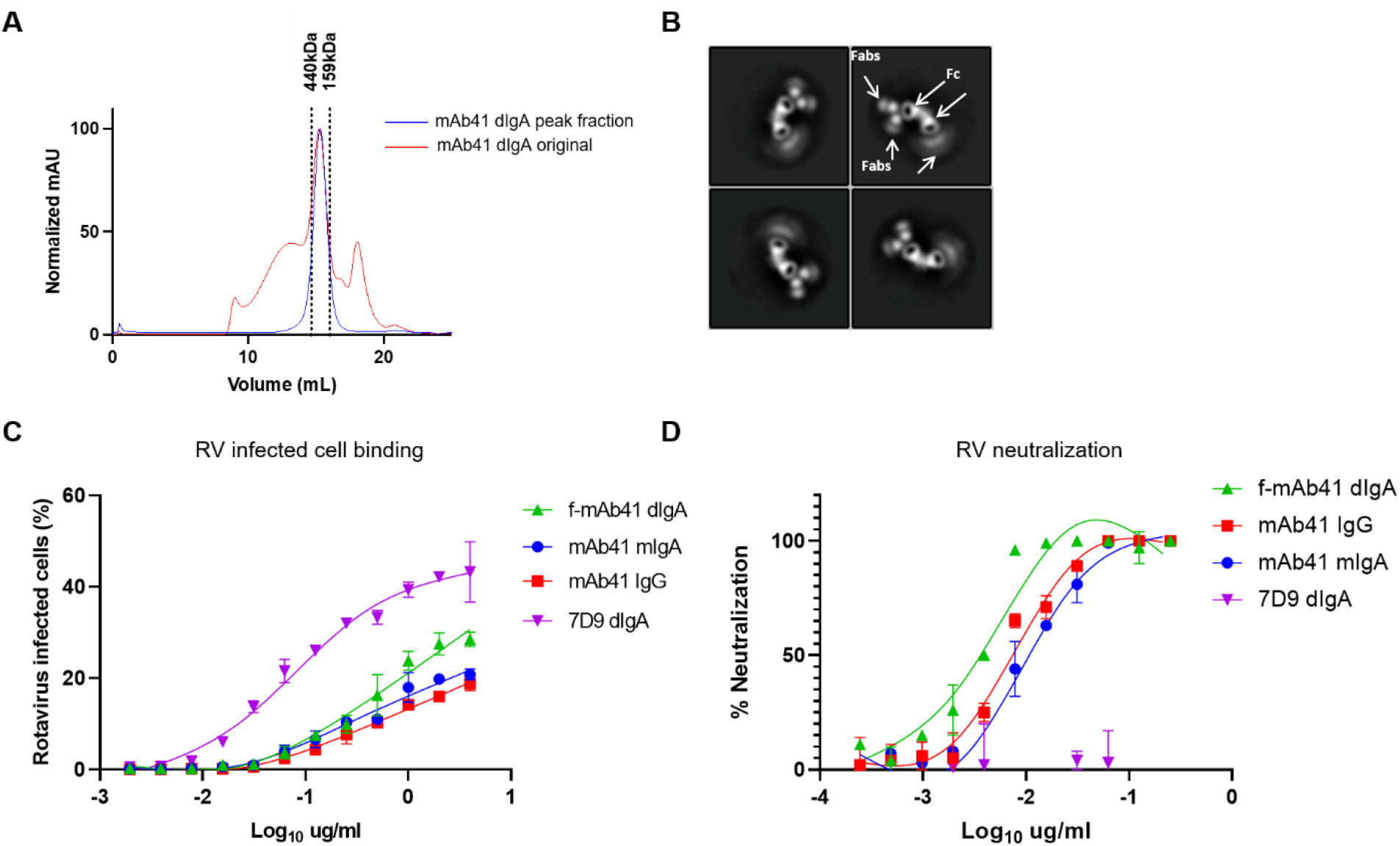
Recombinant mAb41 dimeric IgA antibodies demonstrate greater rotavirus binding and neutralization potency than mAb41 monomeric IgA or IgG antibodies. (A) Size exclusion chromatography by Superose 6 Increase 10/300 GL column in 1XPBS of mAb41 dimeric IgA (dIgA) antibodies at a flow rate of 0.75 ml/min. Molecular weight markers are listed above the dashed lines at 440kDa and 158kDa, respectively. (B) Negative stain electron microscopy of the purified fraction revealed only the presence of dimeric antibodies. Fab and Fc regions are indicated respectively by white arrows. (C) Fractionated mAb41 dimeric IgA (f-mAb41 dIgA) antibodies bound stronger to RV infected cells compared to mAb41 monomeric IgA (mIgA) or IgG as determined by a RV infected cell binding assay. Data are plotted as mean ± SD of replicates. (D) Fractionated mAb41 dimeric IgA (f-mAb41 dIgA) antibodies more potently neutralized RV compared to mAb41 monomeric IgA (mIgA) or IgG as determined by a RV neutralization assay. Data are plotted as mean ± SD of replicates.

### Pharmacokinetics (PK) of intravenous 7D9 dIgA infusion in the blood and milk compartments of lactating BALB/c dams

To determine the optimal dose of dIgA for systemic maternal infusion, we used an Empirical Bayesian estimate of individual PK parameters in both plasma and milk from 7D9 infused dams. Antibody level data were used in simulation for multiple doses including 5 mg/kg (Figure S4A), 10 mg/kg (Figure S4B), and 15 mg/kg (Figure S4C). To estimate the antibody concentrations in serum and milk, 7D9 concentrations were simulated at 24 to 192 hrs in 24-hr intervals. Dosing intervals of 1 to 3 days were explored. Using a 1-day dosing interval, concentrations of 7D9 remained stable up to 8 days after the first dose (Figure S4A). However, with the 2- (Figure S4B) and 3-day (Figure S4C) dosing intervals, 7D9 concentrations dropped by day 2 post-infusion and continued to decrease without an additional dose. The intercompartmental clearance from plasma to milk was 0.11 mL/h and the elimination half-life was 13.55 (6.13 – 18.90) hrs. The observed elimination half-life of 7D9 dIgA is similar to that of dIgA reported in other species (<1 day to ∼4 days), including mice and rhesus macaques (Challacombe and Russell, 1979; Lombana et al., 2019)

### Maternal passive immunization with systemic mAb41 dIgA protects against RV-induced diarrhea in suckling pups

To determine if passive transfer of mAb41 dIgA in milk results in protection from RV induced diarrhea in suckling neonates, we developed a RV challenge model using lactating 129sv mice dams and their pups. We chose 129sv mice strain as 129sv pups develop detectable diarrhea after oral inoculation with human RV (Nair et al., 2017). We first confirmed that similarly to BALB/c mice, both fractionated and unfractionated mAb41 dIgA passively transferred into the milk of 129sv mice after IV infusion (Figure S5). In addition to blood and milk, mAb41 dIgA was also detected in vaginal washes and feces, but not in intestinal content (Figure S5). This was different from what we observed in BALB/c mice (Figure S2), suggesting strain-specific differences in pIgR expression resulting in different rates of passive transfer of the two dimeric IgA antibodies. Next, lactating 129sv dams were injected in the tail vein with 5 mg/kg of f-dIgA mAb41 at 4 to 6 days postpartum (Figure 6A). Due to the short half-life of dIgA in milk, as determined by our PK analysis (Figure S4) and to maximize the amount of f-dIgA mAb41 in the gastrointestinal tract of suckling pups at the time of RV inoculation, pups were inoculated with 1×10^6^ FFU of RV Wa strain between 1 to 2 hrs post dam injection. Litters born to dams injected with 5 mg/kg mAb41 dIgA had lower incidence of diarrhea (7.1%) upon gentle abdomen palpation (Figure 6B,D) compared to litters born to saline-immunized dams (88%) (Figure 6C,D). Significantly lower RV antigen per gram of intestine was observed in pups born to mAb41 dIgA-immunized mothers compared to pups of saline-injected mothers (Figure 6E). Additionally, mAb41 dIgA was detected in the stomach contents of the suckling pups (Figure 6F), which exhibited RV neutralization capacity at the highest dilution measured (Figure 6G) without compromising viability of the cell monolayer (Figure S6). Thus, mAb41 dIgA passively transferred to suckling pups through the milk of lactating dams to protect against RV-induced diarrhea.

**Figure 6.**
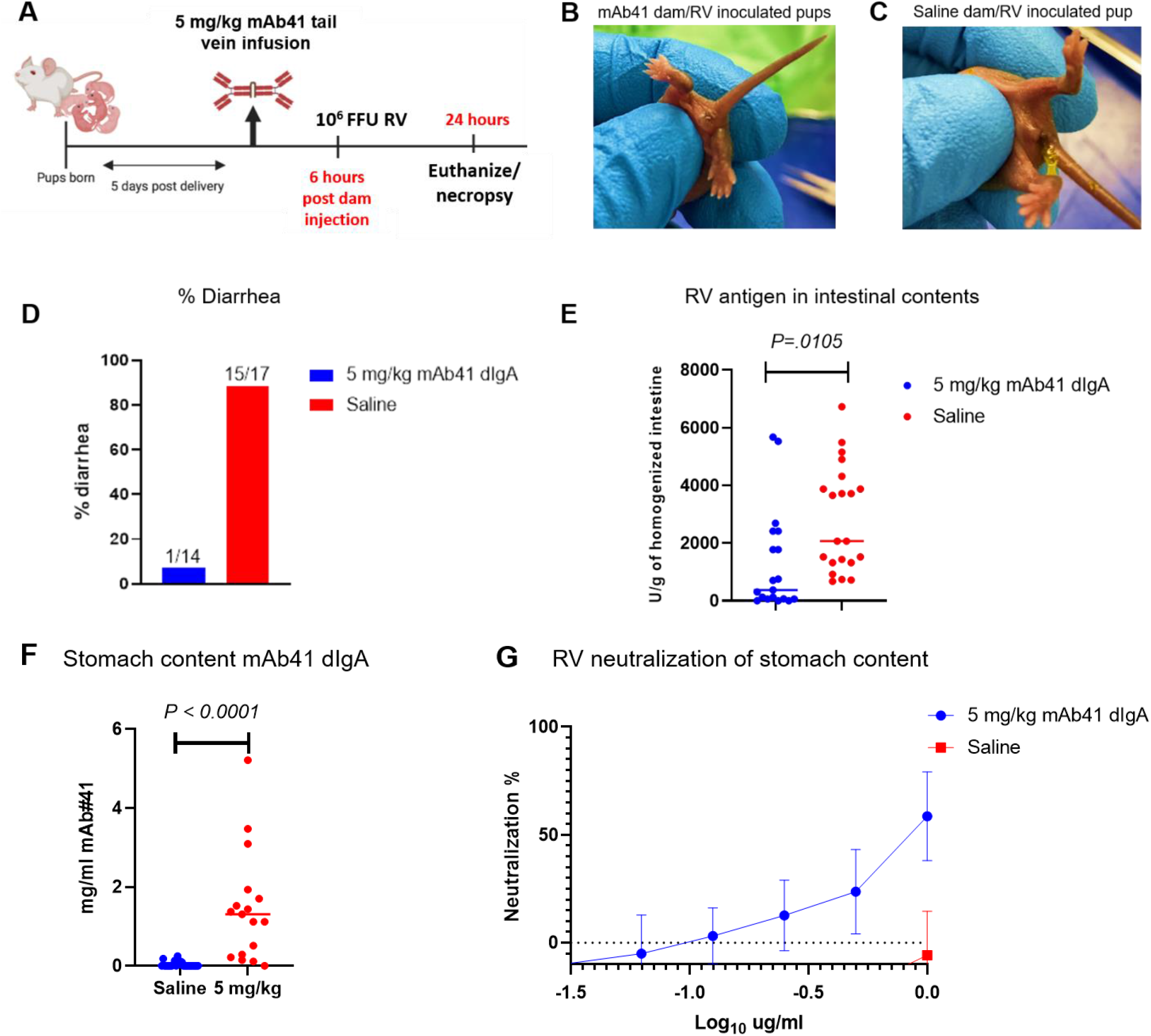
Passive maternal immunization with systemic mAb41 dIgA protects against rotavirus (RV)-induced diarrhea in suckling pups. (A) Schematic of tail vein injections of BALB/c lactating dams with 5mg/kg f-dIgA mAb41 antibodies at 5 days postpartum. Pups were orally inoculated with 1×10^6^ FFU of RV (Wa strain) at 6 hrs post dam injection and euthanized 24 hrs later. Schematic created with Biorender. (B) Representative image of RV inoculated suckling pups from f-dIgA mAb41 infused dams, which excreted urine or hard stool upon abdomen gentle palpation. (C) Representative image of RV inoculated suckling pups from saline infused dams, which excreted yellow, liquid and/or stick stool after gentle abdomen palpation. (D) Diarrhea was reported as % of animals with clinical symptoms upon gentle abdomen palpation in each treatment group (5 mg/kg = blue; saline = red). The number of animals with diarrhea out of the total number of animals are reported at the top of each bar graph. (E) RV antigen in homogenized intestinal tissue was detected via a commercial RV antigen binding ELISA. Data are plotted as individual values for each animal and the horizontal bar represents the median units of RV antigen per gram of homogenized intestinal tissue. Significant differences between the compared groups were determined using a Mann-Whitney U test (**p < 0.05). (F) mAb41 antibodies were detected in stomach content of suckling pups using an mAb41 anti-idiotypic IgA antibody ELISA. Data are plotted as individual values for each animal and the horizontal bar represents the median ng/ml of mAb41 IgA antibodies. Significant differences between the compared groups were determined using a Mann-Whitney U test (***p < 0.001). (G) Stomach content was assessed for rotavirus neutralization at different dilutions and plotted as neutralization % in n=6 pups per treatment group (5 mg/kg = blue; saline = red). Data are plotted as the mean ± SD.

## DISCUSSION

Breast milk contains high levels of sIgA that act as the first line of defense against enteric infection in suckling infants (Glass and Stoll, 1989; Ruiz-Palacios et al., 1990; Torres and Cruz, 1993). Despite this, the burden of RV disease in LMIC remains high, likely due to a combination of factors including high pathogen load, malnutrition and decreased vaccine efficacy compared to high income countries (Guerrant et al., 2008; Otero et al., 2020; Velasquez et al., 2018). Neutralizing antibodies against the external RV proteins VP4 and VP7 play a role in protective immunity against RV infection (Clarke and Desselberger, 2015; Desselberger and Huppertz, 2011; Greenberg et al., 1983; Offit and Blavat, 1986). Therefore, enhancing RV-neutralizing sIgA in breast milk is an ideal strategy to provide protection against RV disease in the suckling infant. However, to date, there are no mAb therapies approved or tested for the treatment or prevention of neonatal infection via passive transfer into breast milk in either humans or animal models. In this study, we sought to develop therapeutic strategies that deliver maternal sIgA into breast milk and could enhance protection against RV disease in the suckling neonate.

We first developed a mouse lactation model and demonstrated that a systemically administered a RV non-neutralizing 7D9 dIgA purified from a previously described hybridoma (Greenberg 1996) passively transfers into dams’ breast milk and the gastrointestinal tract of suckling pups. We then investigated the half-life of a RV neutralizing mAb41 dIgA. After systemic infusion, the elimination half-life was short [13.55 (6.13 – 18.90) hrs] compared to what is reported for circulating IgG (15-30 days depending on the subclass) (Mankarious et al., 1988) and was consistent with previous reports (Lombana et al., 2019). Varying amounts of mAb41 dIgA were also detected in the intestinal content, vaginal washes, and saliva. These data suggest that in addition to breast milk, systemically-administered dIgA can traffic to other pIgR-expressing mucosal sites. Dimeric IgA mAb therapeutic strategies that target a particular site, like the mammary gland, will need to consider the biodistribution into other mucosal tissues. Future work should focus on improving dIgA half-life using viral vectors or nucleic acid delivery systems, and on developing strategies to target specific mucosal tissues.

Exploration of the therapeutic potential of neutralizing dIgA *in vivo* has been hampered by the difficulties in production and purification of dIgA at desired antibody quantities (Reinhart and Kunert, 2015; Virdi et al., 2016). We therefore generated several plasmid constructs to optimize the production of a potent RV-neutralizing dIgA (mAb41). We observed that the construct in which the J-chain gene was placed after the heavy and light chain genes produced the highest level of dIgA following recombinant production. This suggests that spatiotemporal production of the IgA heavy, light and J-chain in the cell influences dimerization and therefore, dIgA production. While there are few studies investigating the molecular mechanisms of J-chain protein production and IgA multimerization *in vitro* or *in vivo*, a recent study demonstrated that the B cell chaperone protein MZB1 plays a role (Xiong et al., 2019). Recombinant production of dIgAs in cells expressing MZB1 may improve dimer formation. We also observed that decreasing the amount of J-chain DNA relative to heavy and light chain DNA, resulted in higher levels of dIgA. These results are not surprising, given that each dimer comprises two IgA monomers linked together by a single J-chain and are likely due to an imbalance in the ratio of available heavy and light chains that can dimerize with J-chain. Functional characterization of the newly generated mAb41 dIgA demonstrated that the dimer had higher neutralization potency compared to both IgA and IgG monomers which may be due to the higher number of available binding sites present on dimers compared to monomers. Isotype-specific mAb protection has been previously demonstrated in both *in vitro* and *in vivo* studies. Mice were better protected from influenza infection in the nasopharynx after systemic administration of anti-influenza polymeric IgA antibodies compared to IgG (Renegar and Small, 1991a, b; Renegar et al., 2004). Interestingly, however recombinantly produced poliovirus-specific antibodies had similar neutralization activity whether they were produced as mIgA, dIgA or IgG (Puligedda et al., 2020) which suggests that isotype-specific differences in functional capacity may be pathogen and even epitope specific.

Finally, using the 129sv mouse model of human RV challenge we showed that a systemically-administered RV-neutralizing dIgA antibody can passively transferred into breast milk and protected suckling neonates from RV-induced diarrhea. Our data support the development of passive immunization strategies with neutralizing dIgA in lactating women to reduce mother-to-child transmission of breast milk-transferred enteric infections including RV, norovirus, and poliovirus, and non-enteric infections like HIV. While passive antibody transfer studies via breast milk have yet to be performed in human infants, oral delivery of recombinant mAbs has been explored. Interestingly, while orally fed palivizumab (anti-RSV IgG1 mAb) was not stable across the infant gastrointestinal tract (Lueangsakulthai et al., 2020a), natural anti-RSV IgG and IgA from breast milk were stable through all phases of simulated infant digestion (Lueangsakulthai et al., 2020b). This demonstrates that delivering pathogen-specific sIgA via breast milk may be a more attractive strategy than oral feeding. The differences in stability between breast milk-derived and recombinant anti-RSV antibodies may be due glycosylation differences. Indeed, recombinant antibodies are differentially glycosylated compared to endogenous antibodies and IgA is more heavily glycosylated than IgG, resulting in altered function (Higel et al., 2016; Langel et al., 2020).

The protective transfer strategy is attractive for clinical translation for several reasons: (1) dIgA antibodies directly traffic to mucosal sites, including the mammary gland and passively transfer into breast milk and then to the infant digestive tract as sIgA; (2) dIgA/sIgA may have increased capacity for virus neutralization compared to IgG; (3) breast milk sIgA is more resistant to proteolysis in the stomach and/or gut compared to other isotypes oral delivery of recombinant sIgA in infants; 4) the short-half-life of dIgA may circumvent maternal anti-drug reactions; and (5) dIgA in circulation will traffic to other maternal mucosal sites including the gut, providing dual protection of the maternal-neonatal dyad against enteric infection. Future work could focus on increasing the half-life of dIgA and developing strategies to target specific mucosal tissues. Development of novel strategies that reduce infant mortality against enteric infections is imperative to reach the World Health Organization’s goal to end preventable deaths of newborns by 2030. Our results will help guide the development of novel maternal immunization strategies, which may leverage passive transfer of neutralizing dIgA into breastmilk and decrease infant morbidity and mortality against enteric pathogens.

## METHODS

### Cells and viruses

African Green Monkey kidney epithelial cell line MA104 (CRL-2378.1) was obtained from American Type Culture Collection (ATCC) and cultured in MEM-alpha (Life Technologies) supplemented with 10% fetal bovine serum (FBS), 50 µ/ml of penicillin and 50 µg/ml of streptomycin (Invitrogen). Rotavirus strain A (Wa) (ATCC) was propagated in MA104 cells as previously described (Patton et al., 2009).

### Rotavirus quantification

RV was quantified using a fluorescence focus forming assay (Patton et al., 2009). In brief, RV was activated with 10 µg/ml of trypsin for 1 hr in a 37°C water bath. Serially diluted virus was added to confluent MA104 cells and incubated for 1 hr at 37°C. Inoculum was removed and growth medium including DMEM (Life Technologies), 5% FBS, 50 µ/ml of penicillin and 50 µg/ml of streptomycin (Invitrogen) was added. Infected cells were then incubated at 37°C for 12 to 18 hrs. Medium was removed from the plates and fixed with 10% formalin in neutral buffered saline for 20 minutes. Wells were then washed with 2% FBS and cells were permeated with 0.5% Triton-X in PBS for an additional 15 minutes. Wells were washed twice and 7D9 (VP6-specific, murine IgA antibody) was added at 10 µg/ml in 2% FBS as the primary detection antibody for 1 hr at room temperature (RT). Cells were washed twice and an anti-murine IgA antibody conjugated to FITC (1:100; Southern Biotec) was added to wells for 1 hr at RT. Cells were washed four times with wash solution and DRAQ5 nuclear stain (Fisher Scientific) was added to cells at 1:2000 dilution. Cells were washed once with PBS and resuspended in 10 µl of PBS. Infection was quantified in each well by automated cell counting software using a Cellomics Arrayscan VTI HCS instrument at ×10 magnification. Subsequently, the percent of infected cells was determined as FITC^+^DRAQ5^+^ cells.

### 7D9 Antibody production

The 7D9 hybridoma line was cultured in ClonaCell™-HY Medium E (STEMCELL Technologies) prior to antibody production. To produce large quantities of 7D9, cells were resuspended in Hybridoma Serum Free Medium (Fisher Scientific) and seeded in the cell compartment of a bioreactor, with Medium E providing nutrients from the medium compartment. Antibody was harvested from the cell supernatant after 5 to 7 days post inoculation and purified using Protein L Sepharose beads (Thermo Fisher Scientific).

### Construction of mAb41 plasmids and antibody production

To generate the human-mouse chimeric mAb41 IgA and IgG, we took the variable domain sequences of a previously isolated human anti-RV VP4 specific neutralizing mAb (*i*.*e*., mAb#41) (Nair et al., 2017), attached it to the murine IgA and IgG constant regions (accession numbers: AB644393.1, JQ048937.1, KT336476.1, JQ048937.1), respectively, and cloned it into the pcDNA3.1 expression vector. A third plasmid encoding the BALB/c J-chain sequence (accession number: AB664392.1) was also generated. The heavy, light and J chain of mAb41 IgA were also cloned into a single open reading frame, and in different orientations as shown in Figure 4. The 2A self-cleaving peptide technology was used to express both the heavy, light and J chain genes from a single open reading frame. Antibodies were produced by transient transfection of human epithelium kidney 293T Lenti-X cells (Clontech Laboratories, Mountain View, CA) using the JetPrime transfection kit (Polyplus Transfection Illkirch, France) following the manufacture’s recommendations. Different amounts of each plasmid were transfected as shown in Figure 4. Antibodies were harvested from cell supernatants at 4 to 5 days post transfection and purified using CaptureSelect™ LC-lambda (mouse) Affinity Matrix (Thermo Fisher Scientific).

### Dimeric IgA characterization and purification

Dimeric antibodies (7D9 and mAb41) were characterized and fractionated by size exclusion chromatography using a Superose 6 10/300 GL on an AKTA liquid chromatography system and concentrated on AmiconUltra 100k spin columns (Millipore).

### Negative-stain electron microscopy (NSEM)

Antibodies were diluted to 100 mg/ml final concentration with buffer containing 10 mM NaCl, 20 mM HEPES buffer, pH 7.4, 5% glycerol and 7.5 mM glutaraldehyde. After 5-minute incubation, excess glutaraldehyde was quenched by adding sufficient 1 M Tris stock for a final Tris 75 mM for 5 mins; then samples were stained with 2% uranyl formate. Images were obtained with a Philips 420 electron microscope operated at 120 kV, at 82,000 × magnification and a 4.02 A° pixel size. RELION 3.0 (Zivanov et al., 2018) was used for CTF correction, automatic particle picking and 2D class averaging of the single-particle images.

### Animals

Timed pregnant BALB/c and 129sv mice were obtained from Charles River laboratories and Taconic Biosciences, respectively. Upon arrival, all mice were maintained in a pathogen-free animal facility under a standard 12 hr light/12 h dark cycle at RT with access to food and water *ad libitum*. Timed pregnant mice received a supplemental nutritional gel to decrease risk of pup savaging. For IV injections of recombinant dIgA mAbs, animals were restrained using a mouse tail vein restrainer. For mouse milking, dams were separated from their pups for at least 2 hrs to allow milk accumulation while pups were kept warm on a heating pad. Dams were administered 2 IU/kg of oxytocin via intraperitoneal (IP) injection. The mammary area was wiped with sterile alcohol prep pad before manually expressing the teat with thumb and forefinger to gently massage the mammary tissue in an upward motion until a visible bead of milk formed at the base of the teat. A sterile pipet tip was used to gently pull the milk into the tip. All teats were milked two times. Milk was diluted 1:4 with PBS and filtered with 0.22 μm Spin-x centrifugal filters (Costar) at 4°C at 15,000 x g for 30 min. The Spin-x filter separated the lipid portion of the milk from the liquid whey portion, and the liquid whey portion was stored in -20°C. Blood samples were collected from the facial vein (submandibular). The blood was allowed to clot at ambient temperature. Clotted blood samples were maintained at RT and centrifuged for 6,000 RPM for 15 min. The serum was separated from the blood and stored at -20°C.

For RV infection, neonatal 129sv mice (5 days old) were orally gavaged with a minimum of 1×10^6^ FFU of RV Wa. Pup stomach contents and intestines were collected and homogenized in 500 µm PBS using a TissueLyzer II (Qiagen) for 5 min at 50 Hz with a stainless steel ball added as a pulverizer. Pulverized stomach content and intestinal tissue were transferred to a new microcentrifuge tube and spun for 10 minutes at 3,000 RPM. Supernatants were collected and then filtered via 0.22-µm Spin-x centrifugal filter tubes by centrifugation at 18,000 × *g* for 20 mins at 4°C. A protease cocktail (1X) (Fisher Scientific) was added (and samples were stored at -20°C until further analysis.

### Biodistribution studies

After IV injection of recombinant mAbs, mice were euthanized via CO_2_ asphyxiation. Mice oral and vaginal cavities were immediately washed with 100 µL of PBS. Oral and vaginal washes were centrifuged at 3,000 RPM for 10 mins to pellet any cellular debris. The supernatant was then collected and stored at -20°C. Additionally, intestinal and rectal contents were collected and diluted in 500 µl of PBS. The diluted samples were then filtered via 0.22-µm Spin-x centrifugal filter tubes by centrifugation at 15,000 x g for 30 mins. Protease inhibitors were added (1X) and samples were stored at -20°C until further analysis.

### VP6 binding IgA antibody ELISA

Recombinant VP6 protein (head domain; residues 147-339 of full-length VP6) was expressed in *E. coli* and purified through affinity chromatography using a Ni-NTA column and size-exclusion chromatography using a Superdex 200 10/300 GL column as previously described (Aiyegbo et al., 2013). Nunc® Maxisorp™ 384-well plates were coated with 3 µg/ml of recombinant VP6 protein diluted in coating solution concentrate (Seracare) overnight at 4°C. Plates were washed one time (PBS, 0.5% Tween-20) and incubated for 2 hrs with blocking solution (PBS, 4% whey protein, 15% goat serum, 0.5% Tween-20). Antibodies were diluted in blocking solution and added to wells in duplicate for 1 hr. Plates were then washed twice and incubated for 1 hr with an HRP-conjugated, goat anti-mouse IgA antibody (Southern Biotech) at a 1:5000 dilution. After 4 washes, SureBlue Reserve TMB Microwell Peroxidase Substrate (KPL) was added to the wells for 10 mins, and the reaction was stopped by addition of 1% HCl solution. Plates were read at 450 nm. OD values within the linear range of a standard curve were used to interpolate the concentration of VP6-binding IgA antibodies in the transfection products. The standard curve was generated by serial dilutions of 7D9 dIgA.

### pIgR binding IgA antibody ELISA

J-chain containing IgA antibodies were measured by pIgR binding ELISA. Nunc® Maxisorp™ 384-well plates were coated with 6 µg/ml of recombinant mouse pIgR protein (R&D Systems) diluted in coating solution concentrate (Seracare) overnight at 4°C. The ELISA assay was completed as described above. OD values within the linear range of a standard curve were used to interpolate the concentration of pIgR-binding IgA antibodies in the transfection products. The standard curve was generated by serial dilutions of 7D9 dIgA.

### mAb41 anti-idiotypic antibody ELISA

Nunc® Maxisorp™ 384-well plates were coated with 1 µg/ml of an mAb41 anti-idiotypic antibody (Biogenes GmbH) diluted in coating solution (Seracare) overnight at 4°C. The ELISA assay was completed as previously described above. OD values within the linear range of a standard curve were used to interpolate the concentration of pIgR-binding IgA antibodies in the transfection products. The standard curve was generated by serial dilutions of mAb41 mIgA antibodies.

### Rotavirus infected cell binding assay

MA104 cells were seeded into 96-well plates and incubated until confluent (3-4 days) at 37°C and 5% CO_2_. RV Wa was thawed at RT and activated with 10 µg/ml of trypsin for 30 mins at 37°C. RV was added to cells at MOI 2 and incubated at 37°C and 5% CO_2_ for 20 to 22 hrs. Cells were fixed with 10% neutral buffered formalin for 20 mins. Cells were washed once with wash solution (2% FBS in PBS). To permeate cell membranes, 0.5% Triton-X in PBS was added to cells for 15 mins. Cells were washed twice and 7D9 added to all wells at 10 µg/ml and incubated for 1 hr in the dark at RT. Cells were washed twice with wash solution and an anti-mouse IgA FITC secondary (Abcam) was added at 1:100 dilution and incubated for 1 hr in the dark at RT. Cells were washed four times with wash solution and DRAQ5 nuclear stain (Fisher Scientific) was added to cells at 1:2000 dilution. Cells were washed once with PBS and resuspended in 10 µl of PBS. Infection was quantified in each well by automated cell counting software using a Cellomics Arrayscan VTI HCS instrument at ×10 magnification. Subsequently, the percent infected cells were determined as FITC^+^DRAQ5^+^ cells.

#### Rotavirus neutralization assay

MA104 cells were seeded into 96-well plates and incubated until confluent (3-4 days) at 37°C and 5% CO_2_. RV Wa was thawed at RT and activated with 10 µg/ml of trypsin for 30 mins at 37°C. Serial dilutions of mAbs or homogenized stomach contents were incubated with RV Wa (MOI = 4) in 50 μl for 1.5 hrs at 37°C. The virus/mAb or virus/plasma dilutions were then added in duplicate to wells containing MA104 cells and incubated at 37°C for 20 to 22 hrs. Cells were fixed with 10% neutral buffered formalin for 20 mins. Cells were washed once with wash solution (2% FBS in PBS). To permeate cell membranes, 0.5% Triton-X in PBS was added to cells for 15 minutes. Cells were washed twice and 7D9 added to all wells at 10 µg/ml and incubated for one hr in the dark at RT. Cells were washed twice with wash solution and anti-mouse IgA FITC secondary (Abcam) was added at 1:100 dilution and incubated for 1 hr in the dark at RT. Cells were washed four times with wash solution and DRAQ5 nuclear stain (Fisher Scientific) was added to cells at 1:2000 dilution. Cells were washed once with PBS and resuspended in 10 µl of PBS. Infection was quantified in each well by automated cell counting software using a Cellomics Arrayscan VTI HCS instrument at ×10 magnification. Subsequently, the ID_50_ was calculated as the sample dilution that caused a 50% reduction in the number of infected cells compared with wells treated with virus only using the Reed and Muench method.

#### Rotavirus antigen ELISA

An EDI fecal rotavirus antigen ELISA kit was used according to the manufacturer’s protocol. In brief, 100 µl aliquot of the homogenized intestinal samples were diluted in kit diluent and added in equal volumes to duplicate wells. A set of standards was included (0, 1.9, 5.6, 16.7, 50, 150 and 300 ng/ml). Samples were incubated for 1 hr at RT. Wells were washed 5 times with washing buffer and incubated with 100 µl of tracer antibody for 30 mins at RT. The wells were washed and 100 µl of the antibody substrate was added. Samples were incubated in the dark for up to 15 mins and 100 µl of stop solution was added to stop the reaction. The absorbance readings were generated at 450 nm. A standard curve was plotted and the antigen concentration in the samples was calculated from the curve.

## Acknowledgements

We thank Bridget Pickle and the Duke’s Division of Laboratory Animal Resources staff for expert assistance. We thank Kathy Yarborough, Jaimie Carter, and Dr. Jamie Peacock at the Duke Protein Production Facility in the Duke Human Vaccine Institute for their assistance in size exclusion chromatography. We thank Dr. So Young Kim in the Functional Genomics Facility for her expertise in the Cellomics Arrayscan VTI HCS instrument. The graphical abstract was created using the BioRender (www.biorender.com). This work was supported by the Bill and Melinda Gates Foundation award OPP1189362 (S.P. and M.B.) and funding from the Translating Duke Health Initiative (P.A.).

## Author contributions

S.N.L. contributed to study design, analyzed the data and wrote the manuscript. S.N.L., J.T., J.C., T.T., H.W., C.E.O., L.W., J.C., H.G. performed experiments, including antibodies production, ELISA, neutralization assays and in vivo studies. W.H. and H.C. performed the pharmacokinetics analysis. R.E. and K.M. performed the negative stain electron microscopy. V.S. and P.A. performed size exclusion chromatography and molecular weight determination. M.B. and S.R.P. conceived the study, oversaw the planning and direction of the project including analysis and interpretation of the data and editing of the manuscript. All authors read, revised, and approved the final manuscript.

