## Supplemental for "Protective Transfer: Maternal passive immunization with a rotavirus-neutralizing dimeric IgA protects against rotavirus disease in suckling neonates"

**Supplemental information**

### Protective transfer: Maternal passive immunization with a systemically-administered rotavirus-neutralizing dimeric IgA protects against rotavirus-induced diarrhea in suckling neonates

Langel SN, Steppe JT, Chang J, Travieso T, Webster H, Otero CE, Williamson LE, Crowe JE Jr, Greenberg H, Wu H, Hornik C, Mansouri K, Edwards RJ, Stalls V, Acharya P, Blasi M, Permar SR

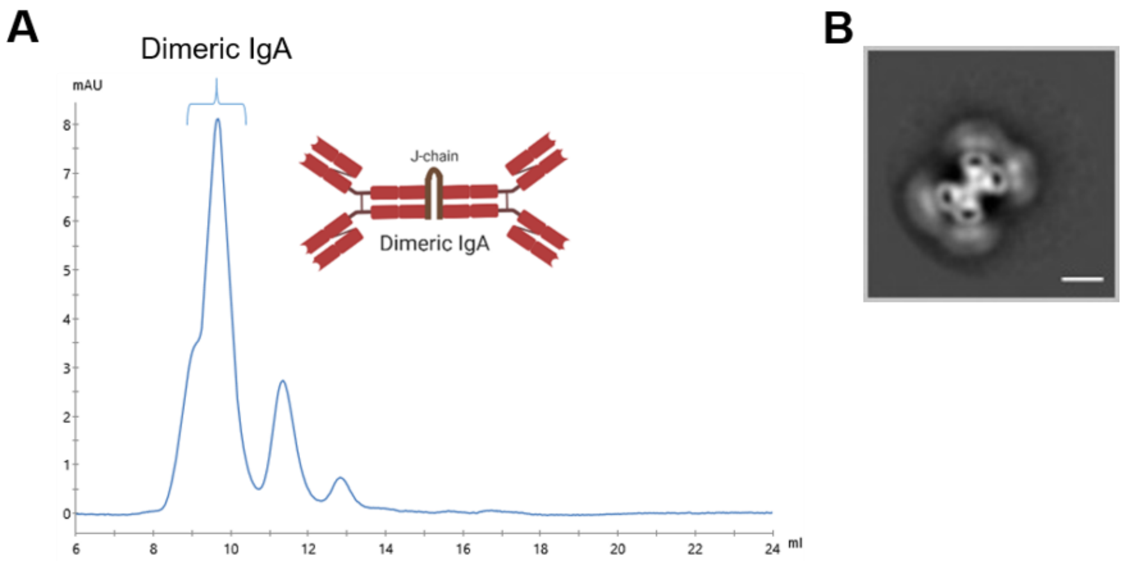


**Figure S1. Characterization of hybridoma-produced 7D9 dIgA**. (A) Size exclusion chromatography demonstrates that most antibodies recovered from the 7D9 hybridoma are dimeric IgA (dIgA). dIgA schematic was created with Biorender. (B) In addition to dimers, higher order IgA species like tetramers are observed via negative stain electron microscopy as shown by the 2D class average. Scale bar represents 10 nm.

A

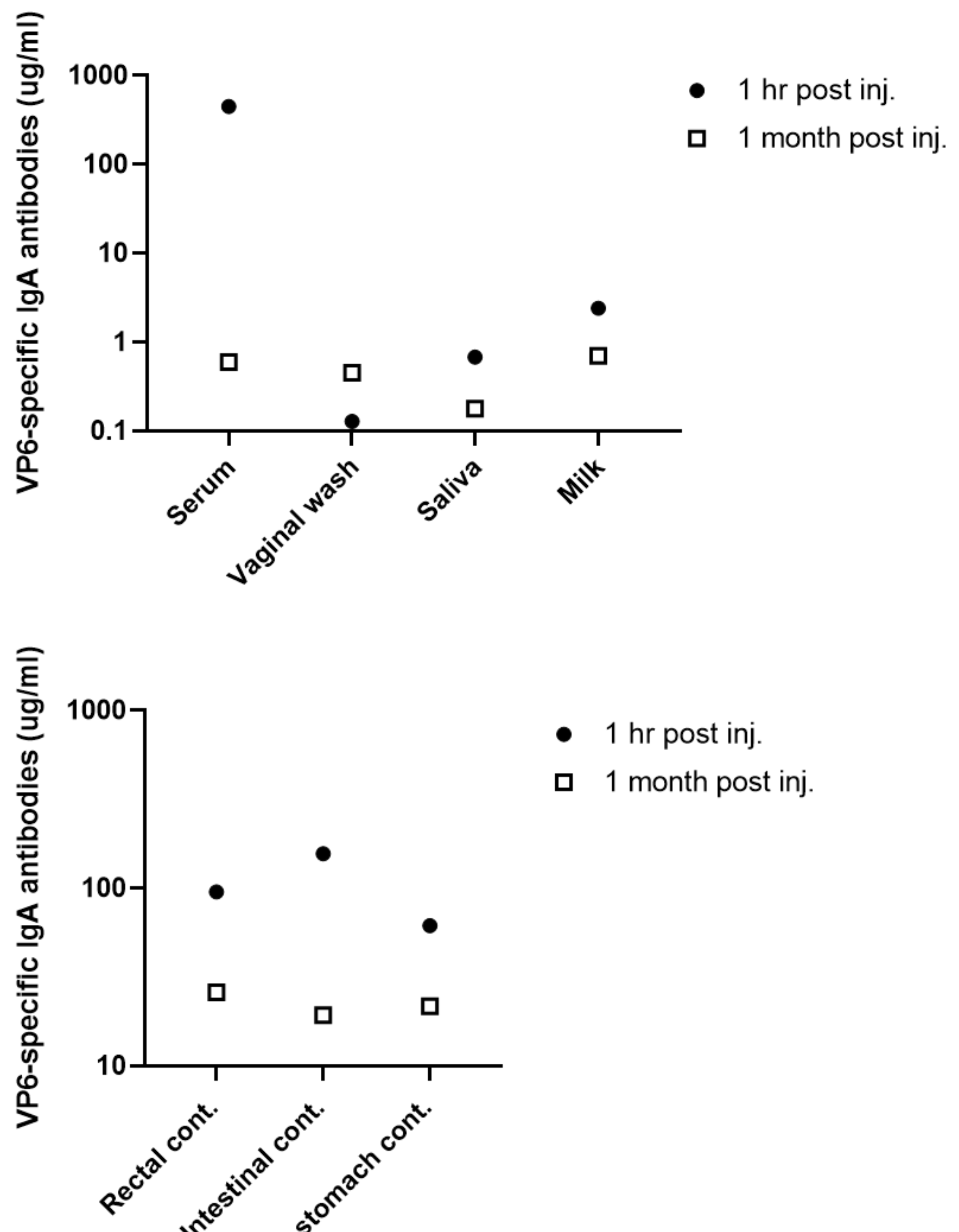


**Figure S3. Biodistribution of tail vein infused 7D9-produced VP6-specific IgA into lactating BALB/c mice and suckling pup stomach contents.** 15 mg/kg 7D9 antibodies were tail vein infused into lactating BALB/c mice (n=2). One lactating dam and one suckling pup were sacrificed at 1 hr post injection (black filled circles). The second lactating dam and an additional suckling pup were sacrificed at 1-month post injection (open squares). The presence of VP6-specific IgA antibodies was analyzed in serum, vaginal wash, saliva, and milk (A) as well as rectal and intestinal content (B) collected at 1 hr and 1 month post-injection via ELISA.

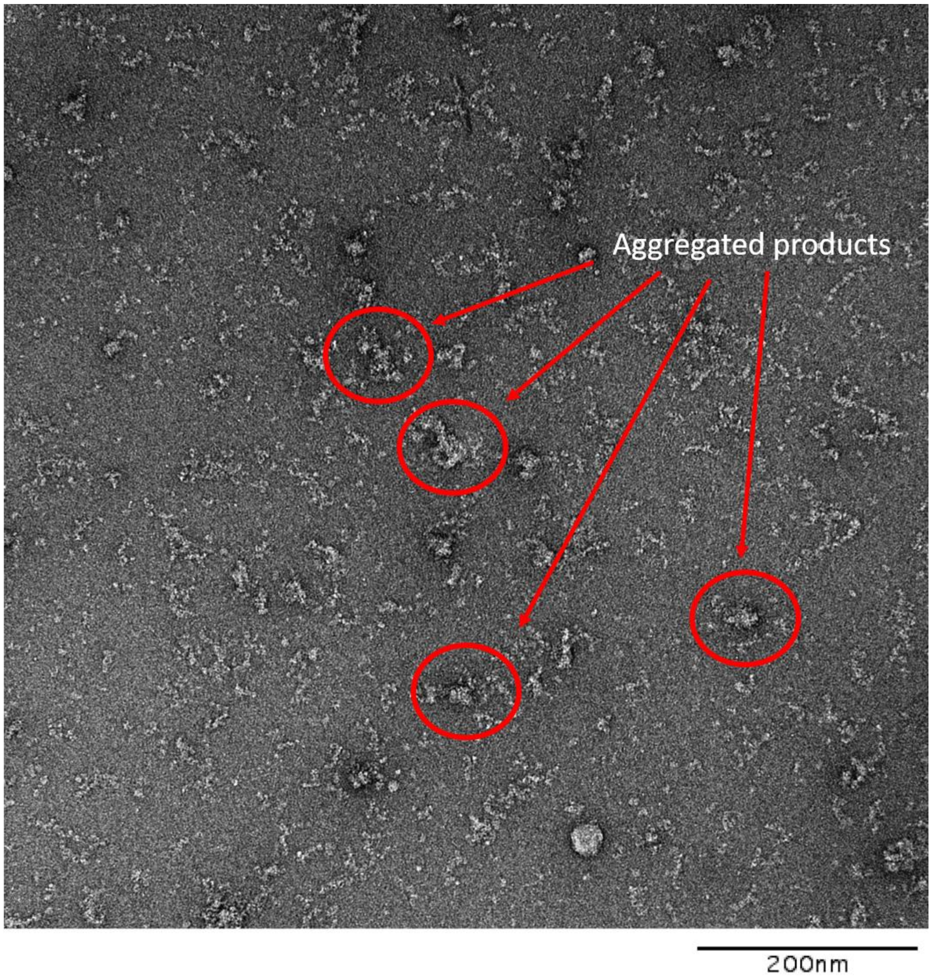


**Figure S3. A representative negative stain electron microscopy image of mAb41 transfection product.** Red circles depict aggregated antibody products. Scale bar is 200 nm.

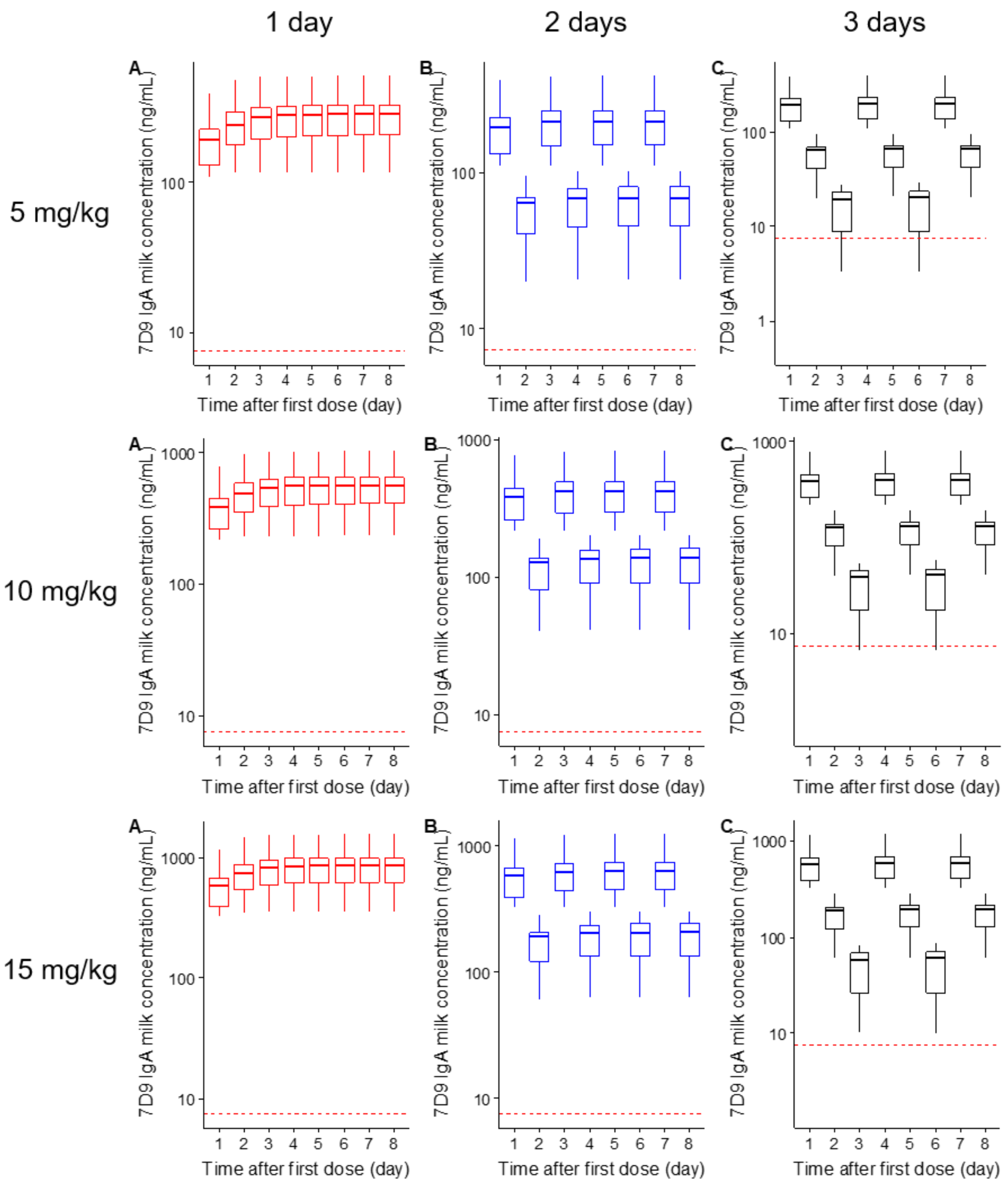


**Figure S4 Pharmacokinetic (PK) analysis antibody levels in periphery and milk following intravenous 7D9 dIgA infusion.** (A) Empirical Bayesian estimates of individual PK parameters of blood and milk from 7D9 infused dams (main text Figure 1D) were used to simulate exposures following various doses of 7D9 IgA. Using a 1-day dosing interval, concentrations of 7D9 dIgA remained stable for up to 8 days after the first dose (A). However, with the 2- (B) and 3-day (C) dosing intervals, 7D9 dIgA concentrations dropped by day 2 post-infusion and continued to decrease without an additional dose.

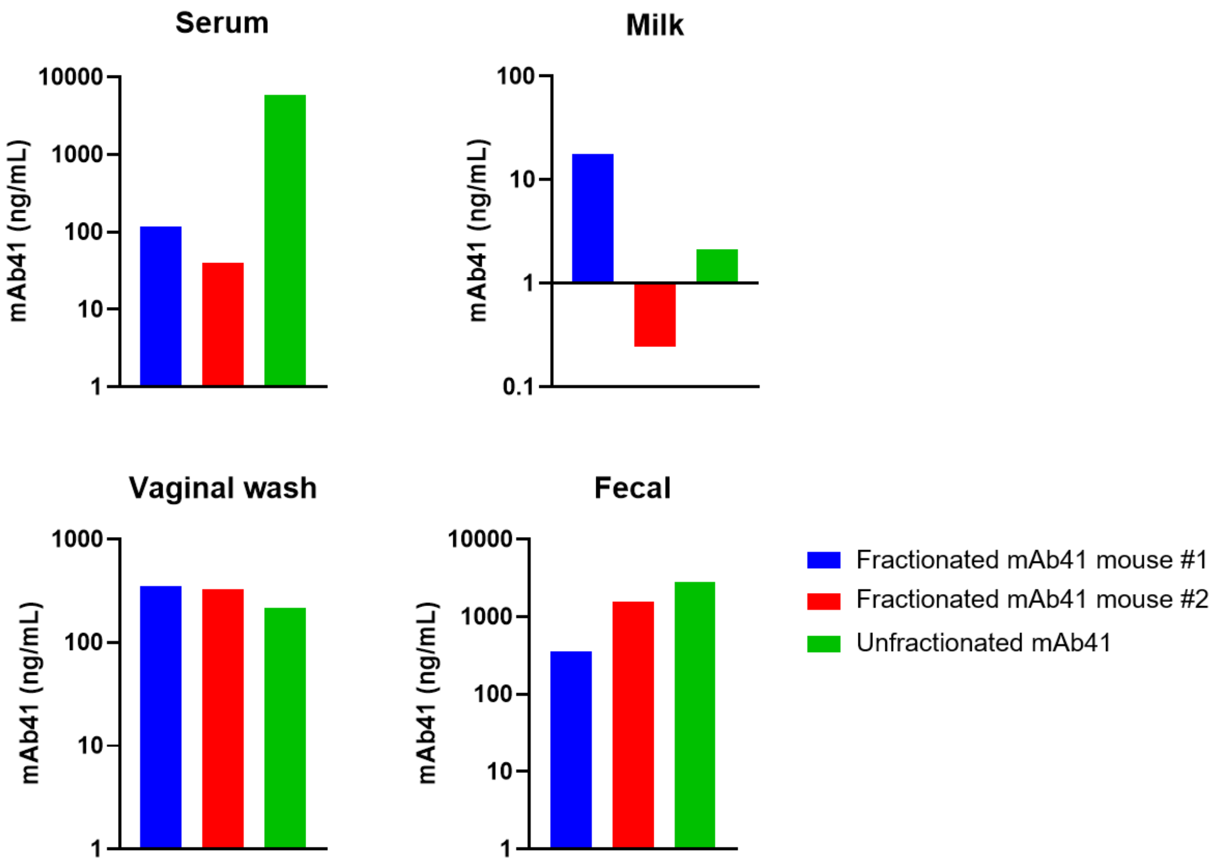


**Figure S5. mAb41 dIgA traffics to mucosal compartments following systemic infusion in 129sv mice.** Two 129sv mice were infused via tail vein injection with 5 mg/kg of fractionated (dIgA enriched) (n=2) antibody. An additional mouse was infused with unfractionated mAb41 IgA. At 2 hrs post-infusion, mAb41 IgA was found in the serum (A), filtered whey (B), vaginal secretions (C), and fecal pellets (D) of the infused mice. Intestinal contents, saliva, nasal secretions, and bronchioalveolar lavage (BAL) fluid were also examined. No mAb41 IgA was present in the intestinal contents or saliva. Data represent mean of assay duplicates.

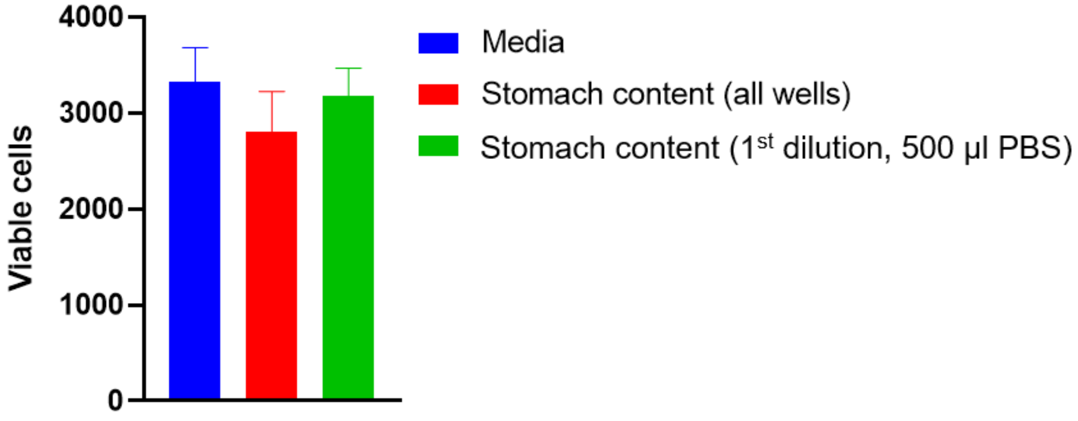


**Figure S6. MA104 cell monolayer viability is not compromised by diluted stomach content samples.** There were no significant differences in MA104 cell viability between wells incubated with diluted (in 500 µl PBS) stomach content compared to medium in the RV infected cell binding assay.
